# Initial data release and announcement of the Fish10K: Fish 10,000 Genomes Project

**DOI:** 10.1101/787028

**Authors:** Guanngyi Fan, Yue Song, Xiaoyun Huang, Liandong Yang, Suyu Zhang, Mengqi Zhang, Xianwei Yang, Yue Chang, He Zhang, Yongxin Li, Shanshan Liu, Lili Yu, Inge Seim, Chenguang Feng, Wen Wang, Kun Wang, Jing Wang, Xun Xu, Huanming Yang, Nansheng Chen, Xin Liu, Shunping He

## Abstract

With more than 30,000 species, fish are the largest and most ancient vertebrate group. Despite their critical roles in many ecosystems and human society, fish genomics lags behind work on birds and mammals. This severely limits our understanding of evolution and hinders progress on the conservation and sustainable utilization of fish. Here, we announce the Fish10K project, an international collaborative project or initiative? aiming to sequence 10,000 representative fish genomes under a systematic context within ten years, and officially welcome collaborators to join this effort. As a step towards this goal, we herein describe a feasible workflow for the procurement and storage of biospecimens, and sequencing and assembly strategies. To illustrate, we present the genomes of ten fish species from a cohort of 93 species chosen for technology development.

## Background

### Fish genomes sequenced to date

As of writing, genome assemblies are publicly available for less than 1% of fish species (216 species of 56 orders) (**Supplementary Table 1**). Their assembly length ranges from 302.36 Mb (*Diretmus argenteus*) to 3.81Gb (*Zeus faber*), with an average length of 813.74 Mb. The average scaffold N50 and contig N50 values are 8.53 Mb and 860.51 Kb, while the median scaffold N50 and contig N50 are 406.7Kb and 20.3Kb, respectively. There are 97 species with a scaffold N50 of more than 1Mb, of which 31 have a contig N50 above 1Mb (**Figure 1**). These genomes has fueled a number of studies on the phylogeny and evolution of fish (e.g., the African coelacanth genome and tetrapod evolution), evolutionary processes of specific fish subgroups (e.g., elephant shark genome illustrating the phylogenetic relationship of Chondrichthyes as a sister group to bony vertebrates) [1], genetic mechanisms of adaptation to different environments (e.g. the deep-sea Mariana Trench snailfish and cave-dwelling fish) [2], and specific biological processes (for example, the tonguefish *Cynoglossus semilaevis* genome for understanding ZW sex chromosome evolution) [3]. Nevertheless, the current fish genome sequencing results are only a drop in the ocean, and numerous critical research questions remain to be resolved. A non-exhaustive list includes gaining a comprehensive and clear understanding of fish phylogeny, genome size diversity and chromosome evolution, diverse environmental adaptations, morphology evolution, respiratory system, immune system, the evolution and function of ultra-conservative (UCE) and conserved nonexonic elements (CNEEs).

**Figure 1.**
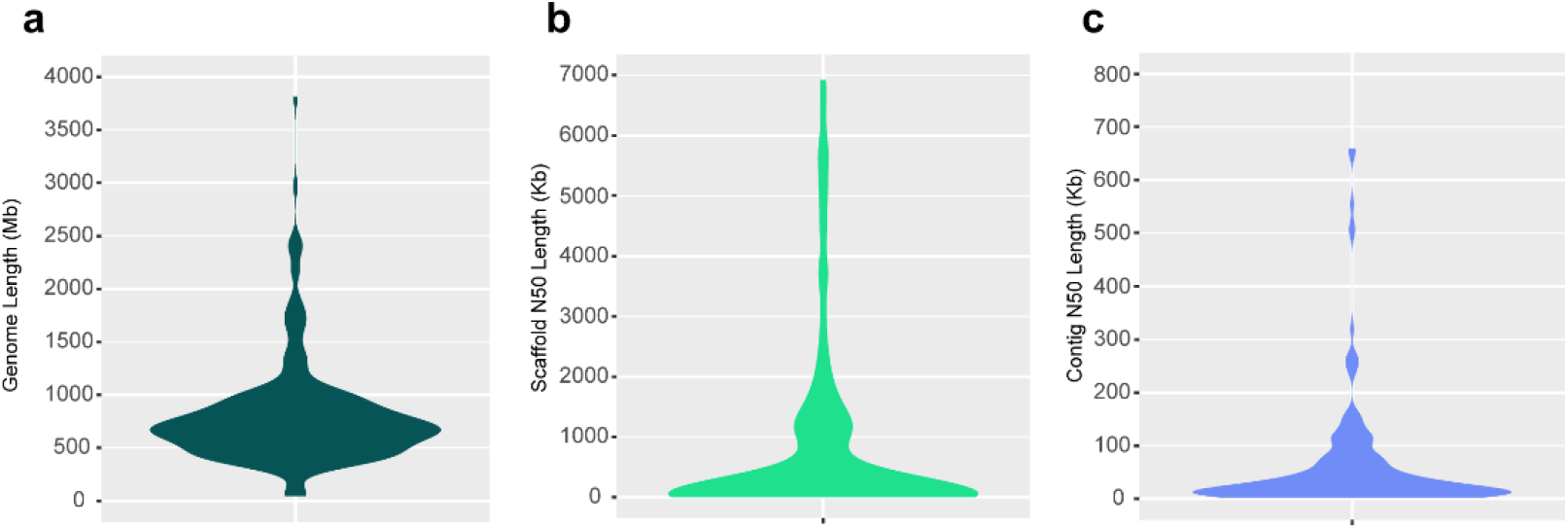
Assembly statistics of fish genomes in public databases. a) Summary of genome size. b, c) N50 statistics. A scaffold is a series of contigs that can be linked. N50 is the sequence length of the shortest contig (or contig) at 50% of the total genome length.

### The era of genome consortiums

With the rapid development of DNA sequencing technology, this is the time for large-scale, collaborative genomic studies. The first such project was the Genome 10K (G10K) Project established in 2009, which aimed to sequence and assemble genomes of about 10,000 vertebrate species [4]. Further advances in sequencing have extended this vision. The Vertebrate Genomes Project (VGP) was launched in 2017 to generate chromosome-level, haplotype-phased genome assemblies of vertebrate species [5]. The Bird 10,000 Genomes Project (B10K) was initiated [6] after the successful phylogenomic study on 45 avian genomes in 2014 [7]. The B10K projects aims to sequence and assemble all known bird species in three phases. Similar efforts have been made for bats [8], plants [9], and other species [10, 11]. Despite current challenges in funding, sampling, sequencing, assembly, and data analysis, these projects have already made substantial progress. For fish, which makes up more than half of all vertebrate species, no projects at the similar scale has been initiated. The only large-scale genomic study to our knowledge was Fish-T1K, which aimed to sequence the transcriptomes (RNA-seq) of ray-finned fishes [12]. However, the insights gained from transcriptome data alone is relatively limited. Accelerating fish genomics by large-scale genome sequencing efforts would undoubtedly boost research into fish biodiversity, speciation, adaptation, as well as aiding the conservation and sustainable utilization of fish.

### The Fish10K Genome Project

We here announce the Fish10K Genome Project, aiming to sample, sequence, assemble, and analyze genomes of 10,000 fish species. We are proposing an effective and integrated workflow, in which major genomics challenges are addressed, to construct high-quality reference genomes. Through developing and applying effective analysis methods, we will be able to address critical evolutionary and biological research questions related to fish. In order to prove the efficiency of our workflow and the feasibility of this large-scale genome project, we are releasing ten high-quality genomes as part of a pilot project. We hope the released genomes, along with the other genomes generated by Fish10K, will be valuable resources for fish researchers as well as to fishery industry.

## Main text

### Feasibility test and the release of ten fish genomes

In order to establish cost-effective strategies and assess the feasibility of a large-scale genome project, we initiated a pilot study in June 2017. Over the last two years, we went on four expeditions across lakes, rivers, and coastal waters of China, collecting 324 fish species. After careful documentation of sample information and species identification, the tissues of 93 species were selected for DNA extraction and sequencing. We used single tube long fragment reads technology (stLFR) [13] and the DNBSEQ platform to sequence the species, generating long read (Nanopore or PacBio) and Hi-C data for a subset. In this way, we were able to test the feasibility of three different sequencing and assembly strategies (**Figure 2**): stLFR data alone (synthetic long reads generated using second-generation sequencing platform) (Strategy I); stLFR data combined with low-depth, long reads (~10× raw Nanopore data to fill in the gaps) (Strategy II); and high-depth long reads (~80× raw Nanopore data) combined with second generation short reads (either short insert size libraries or stLFR) (Strategy III). We have sequenced all the 93 species using stLFR (**Supplementary Table 2**), 26 species using Nanopore at low depth, and 13 using Nanopore at high depth. To date, we have assembled the genomes of 48 species, which are currently undergoing curation, and are finishing assembly for the remaining 57. For the 48 assembled fish genomes, the average contig N50 and the average scaffold N50 are 1.53 Mb and 5.20 Mb, respectively. The average Benchmarking Universal Single-Copy Orthologs (BUSCO) [14] completeness estimate is 93.48% **Supplementary Table 3**). A comparison of assembly statistics revealed that assemblies generated with Strategy II and Strategy III were more continuous, achieving an average contig N50 of 3.99 Mb and an average scaffold N50 of 9.51 Mb. They are notably superior to Strategy I (average contig N50 of 79.04 Kb and average scaffold N50 of 4.73Mb). To illustrate our effort, we are releasing the genomes of ten representative bony fish genomes covering the three assembly strategies (**Table 1**). The contig N50s of seven of these genomes are more than 1 Mb and a minimum 93% of BUSCO genes were found, indicating the genome assemblies are of high quality. Three genomes were assembled at chromosome-level, with more than 92% scaffold-anchored ratio using Hi-C data.

**Table 1.**
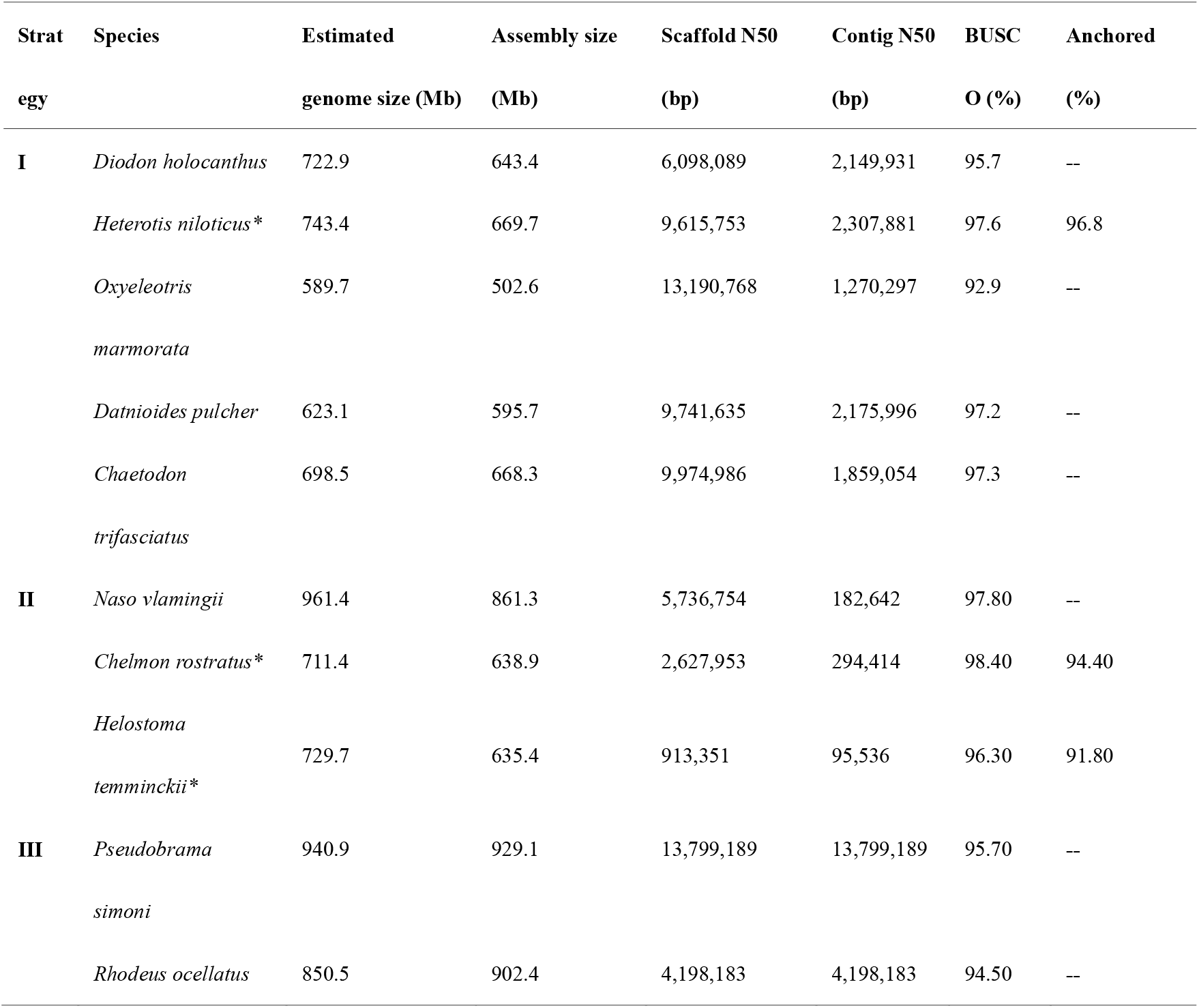
Assembly statistics of the ten released genome assemblies. An asterisk (*) denotes a chromosome-level genome assembly (Hi-C data generated).

**Table 2.**
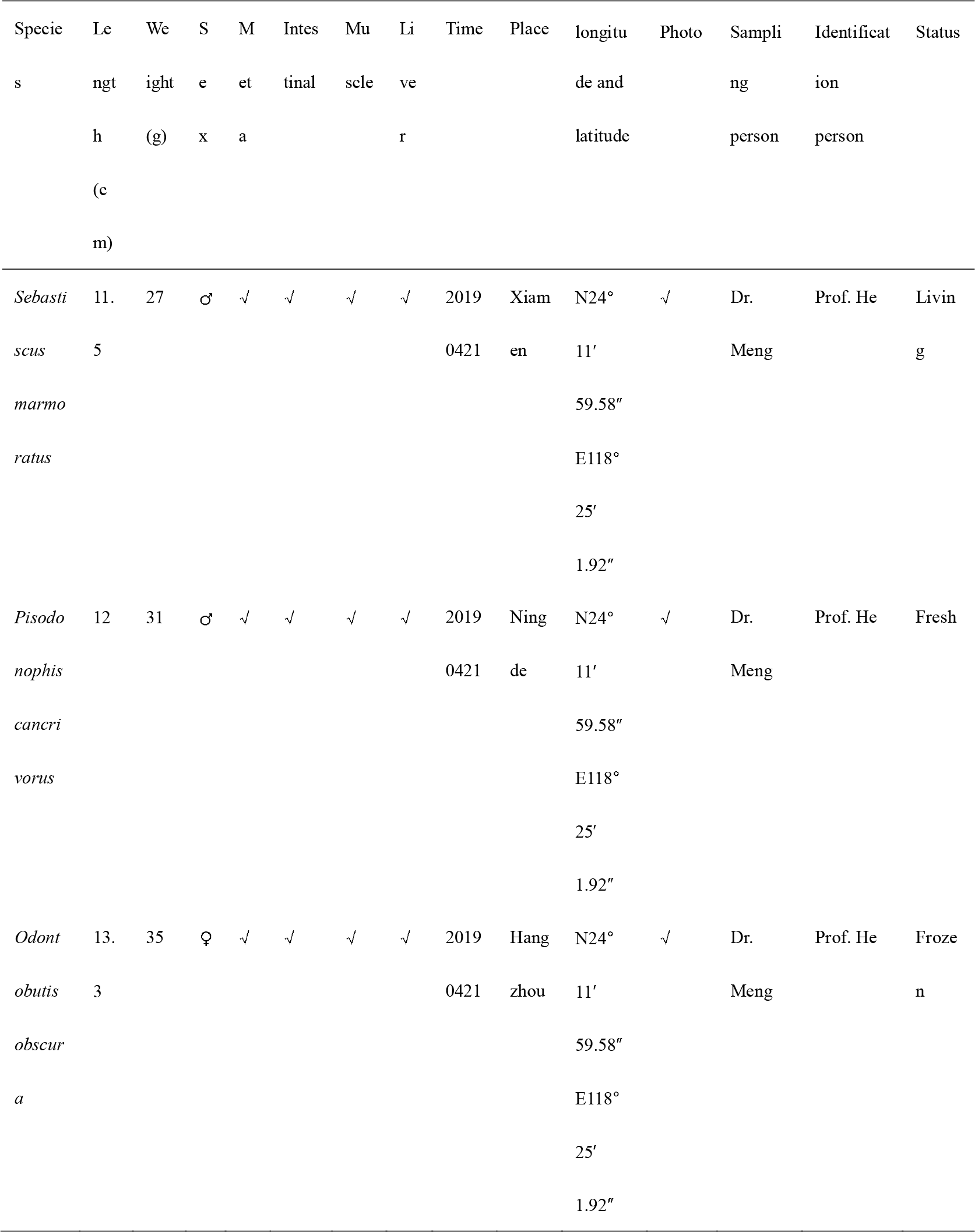
Sample collection template.

**Figure 2.**
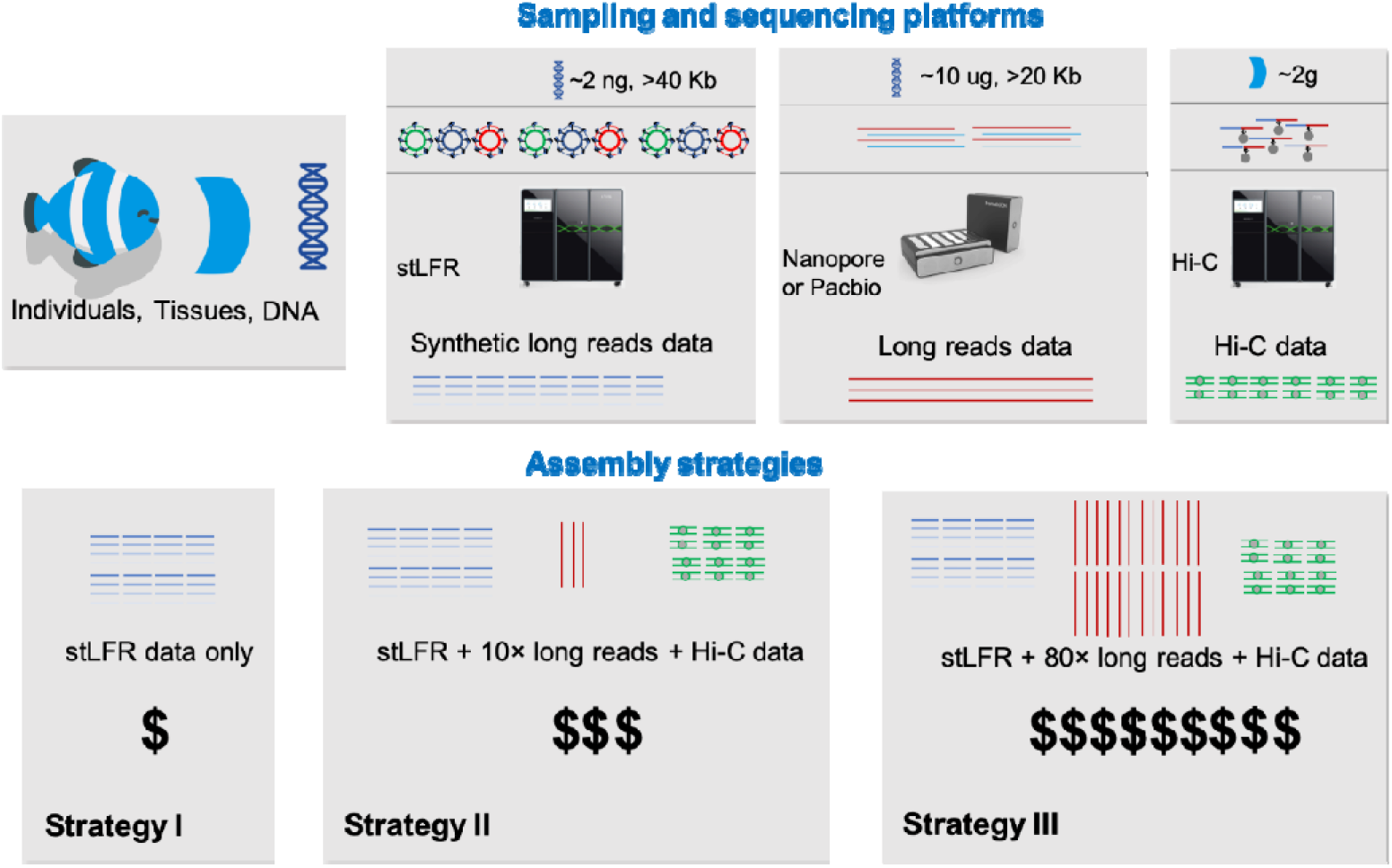
The sequencing and assembly strategies. In the preferred strategy (Strategy II), high-quality DNA fragments (≥40Kb) are used to construct a stLFR library which is sequenced using the DNBSEQ platform. Low-sequencing-depth long reads are only used to improve the continuity of highly complex regions (increase the contig N50). In the alternative Strategy I, high-depth long reads are used to construct contigs, while low-depth stLFR reads are used to polish the contig and link the scaffolds. Hi-C data is used to generate a chromosome-level assembly.

### The Fish10K Genome Project: from 100 to 10,000

With the experience gained in the Fish10K pilot study and our published results, we believe that the project can scale-up. Thus, we are proposing a roadmap (**Figure 3**) in which we will construct high-quality reference genomes for representative species in all orders (Phase I) and families (Phase III) in concert with the generation of draft genome sequences for additional related species (Phase III).

**Figure 3.**
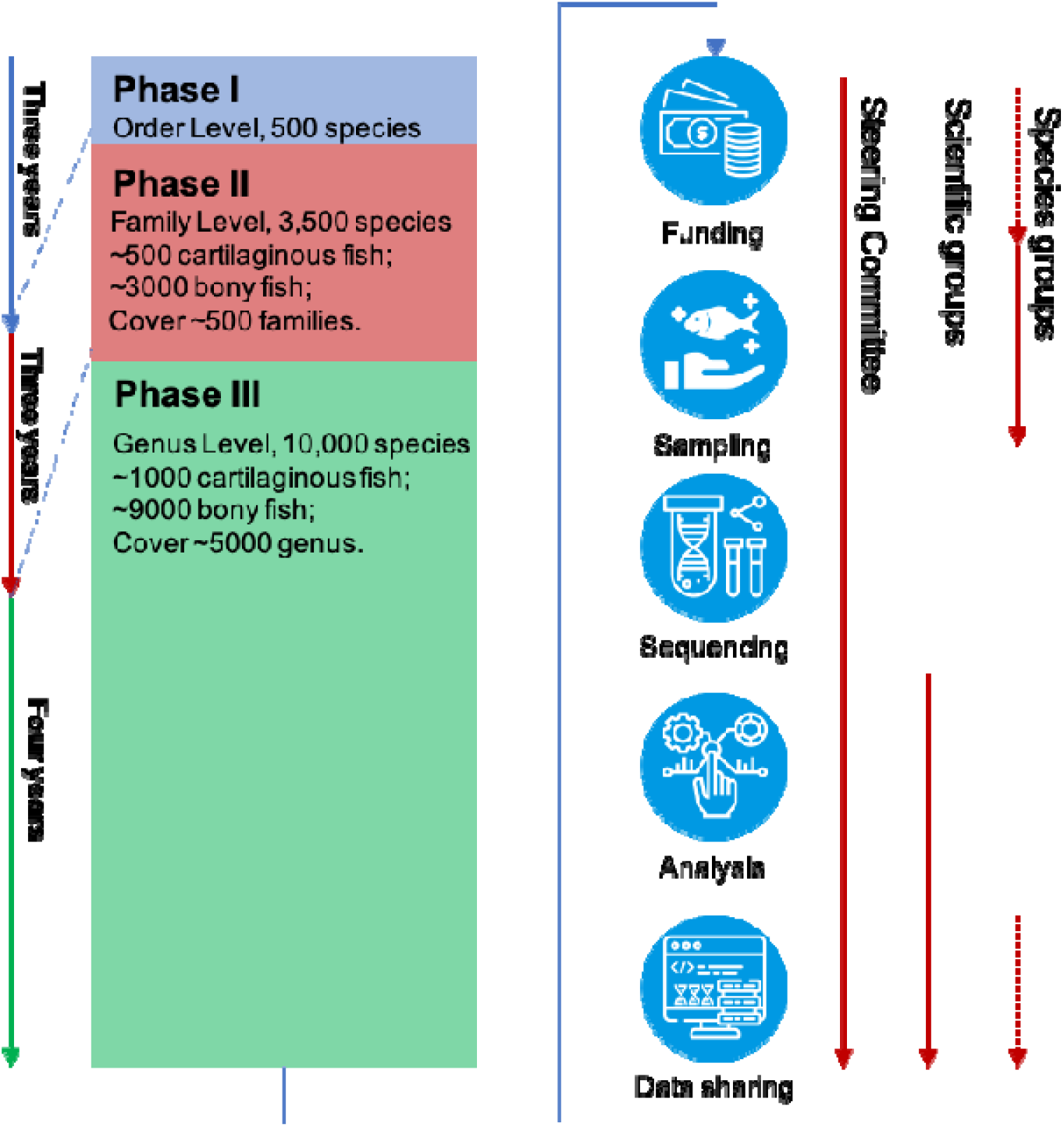
The roadmap and organisation of Fish10K. Fish10K is divided into three phases, based on the evolutionary relationship of fish, and three working groups (steering committee, scientific groups, and species groups).

An interrogation of FishBase [15] and “Fishes of the world” (5^th^ ed.) [16] revealed information on 34,115 fish species from ~5,000 genera, ~529 families, and ~80 orders (**Supplementary Table 2**). The species were divided into six lineages (Elasmobranchii, Holocephali, Actinopterygii, Sarcopterygii, Cephalaspidomorphi, and Myxini), in which Elasmobranchii and Holocephali belong to Chondrichthyes (cartilaginous fishes); and Actinopterygii and Sarcopterygii belong to Osteichthyes (bony fishes). As mentioned above, there are reference genomes available for at least one species of 56 orders, while for the rest of the orders reference genomes are required. Also, there are fish orders with a large number of species (e.g., Perciformes has 62 families; Siluriformes has 40 families; and Scorpaeniformes has 39 families), suggesting that additional high-quality reference genomes are required to represent the diverse biological characteristics. Thus, in Phase I we aim to sequence 450 bony fish and 50 cartilages fish species, covering all 80 orders (**Supplementary Table 2**). In Phase II, we aim to sequence approximately 3,000 species, covering almost all ~500 fish families. In Phase III, we will sequence ~6,500 fish genomes, covering ~5,000 genera.

### Sampling, sequencing, assembly, and annotation

Sampling is a critical challenge in any large-scale genome consortium. We propose a centralised sampling mode (i.e., mirroring our 93-species pilot phase), with several sampling centres set up to collect samples. In addition to these sampling centres, we would like to obtain further samples from around the world. To make sure we have enough information for further analysis and to maximise the value of the genome data, we propose a sampling standard. The associated meta-data was designed to include as much information as possiblestressing the importance of collecting images of each specimen and adequate storage conditions (frozen or voucher specimen).

For sequencing, we propose to use both second- and third-generation sequencing technologies to generate high-quality genome assemblies. Based on our pilot study, and considering the feasibility of obtaining the required amount of high molecular DNA, we have chosen a ‘stLFR data + low-depth Nanopore data + Hi-C data’ strategy (Strategy II in **Figure 2**) for the majority of the species. For more complex genomes, we will generate high-depth Nanopore sequence data to ensure that good assemblies can be achieved (Strategy III, ‘stLFR data + high-depth Nanopore data + Hi-C data’; **Figure 2**). For key species (to be determined by the working groups; see below), we will employ a Pacific Biosciences circular consensus sequencing (CCS) long high-fidelity (HiFi) approach, allowing the generation of highly-accurate long reads [17]. For the large-scale sequencing of 6,000 species in Phase III, we propose to employ stLFR alone (Strategy I in **Figure 2**). For a diploid species with a genome size of less than 10Gb (generated using our preferred Strategy II), we will require the contig N50 and scaffold N50 to be longer than 1Mb and 10Mb, respectively, and (if applicable) to anchor more than 90% of the assembled sequences to chromosomes. The same criteria will apply to assemblies generated using a high-depth long read strategy (Strategy III). For assemblies generated using stLFR sequencing alone (Phase III) assemblies must have a contig N50 and scaffold N50 longer than 100 Kb and 1Mb, respectively. All assemblies must have a BUSCO completeness estimate higher than 90%. Finally, genome feature annotations (e.g., repeat and gene annotations) will be performed using well-established in-house pipelines.

### Fish10k data sharing

Per the Fort Lauderdale [18] and Toronto International Data Release Workshop guidelines [19], all sequencing data (including raw data, assemblies, and annotations) will be deposited in the appropriate INSDC databases, alongside GigaDB and China National Gene Bank (CNGB). The website of Fish10K (http://icg-ocean.genomics.cn/index.php/fish10kintroduction) will provide detailed information on the project status, as well as continuously updated information on the sequenced species. It also provides a portal for data download (in particular for assembled genomes).

### Organisation of Fish10K consortium

Fish10K has been initiated by a core group of researchers, forming the steering committee of Fish10K (**Figure 4**). The steering committee oversees the project and is responsible for fundraising, expanding the steering committee, organising the scientific groups and species groups, and coordinating sampling, sequencing, assembly, and analysis strategies. The steering committee is also responsible for the generation of genomic data. Various scientific groups will focus on technical and scientific questions related to this project. The scientific groups, which will have advance access to all generated data, will include a sampling group, a sequencing and assembly group, and a series of groups focusing on different fish-related scientific questions. We wish to receive proposals from researchers who would like to take part in scientific groups. We also invite researchers who are studying fish species which are rare or extinct to join Fish10K as members in the species group (with or without associated funding for sequencing). In addition to obtaining the genome sequences of their area of interest, joining the consortium provides immediate access to all genomes currently being assembled by Fish10K.

**Figure 4.**
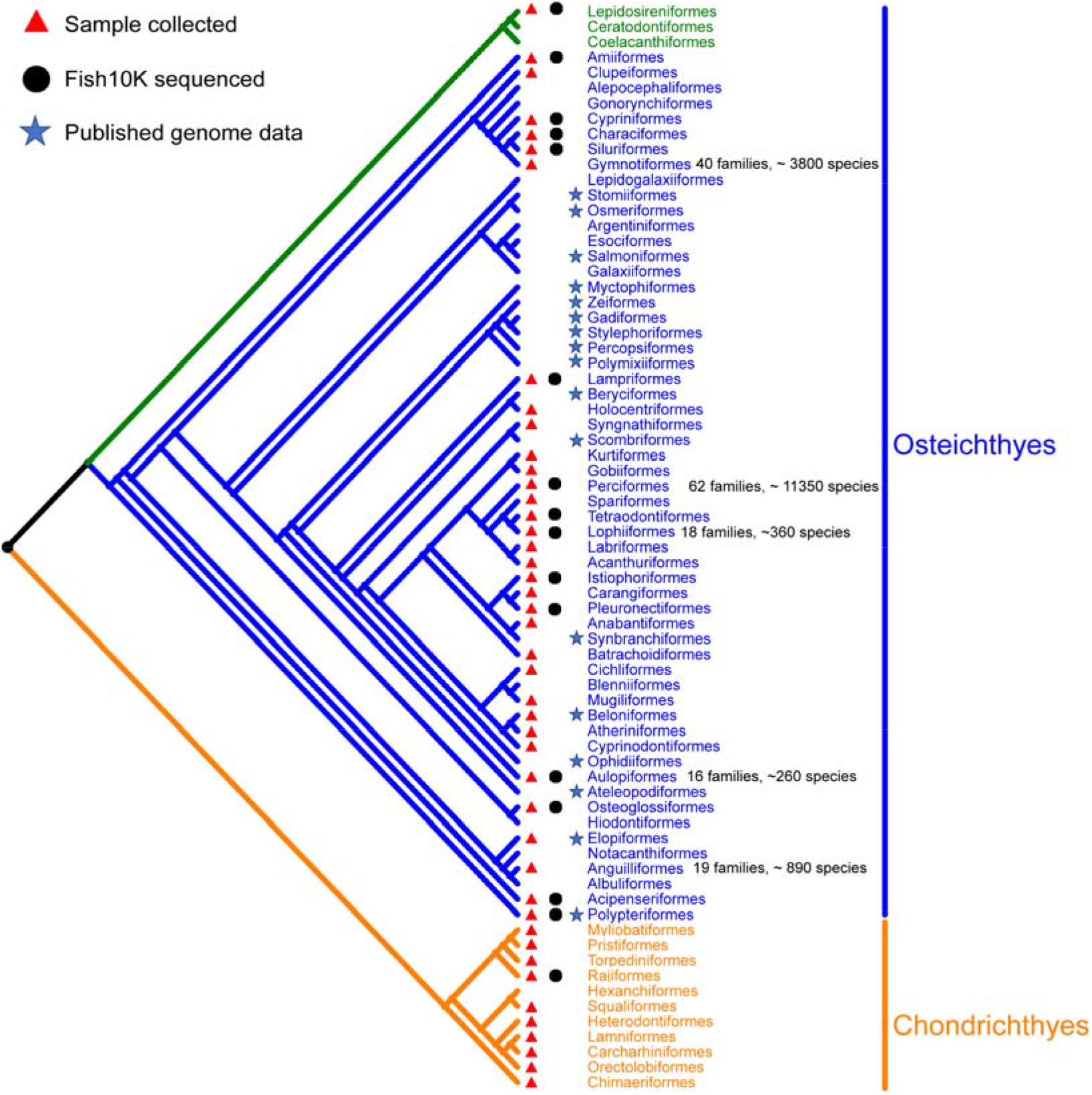
Phylogenetics tree of fish. Jawed vertebrates (gnathostomes) are divided into two major groups, cartilaginous fishes (Chondrichthyes; in orange) and bony vertebrates (Osteichthyes; in blue and green). Bony fishes are grouped into two subgroups (Sarcopterygii; green) and (Actinopterygii; blue). The number of families and species of the five largest orders are labeled. The remaining ten orders of bony fish (Caproiformes, Callionymiformes, Gobiesociformes, Icosteiformes, Lepisosteiformes, Moroniformes, Scombrolabraciformes, Scorpaeniformes, Trachichthyiformes and Trachiniformes) and two orders of cartilaginous fish (Rhinopristiformes and Squatiniformes) are not included in the phylogenetic tree, due to their uncertain position.

## Conclusions

Fish10K will generate an unprecedented, comprehensive data set of fish, the largest and most diverse vertebrate group. Our effort will allow us to complete the genomic tree for fish and, in concert with other projects such as VGP and B10K, vertebrates in general.

## Abbreviations

B10K: Bird 10,000 Genomes Project
VGP: Vertebrate Genomes Project
EBP: Earth BioGenome Project
Fish-T1K: Transcriptomes of 1,000 Fishes Project
Fish10K: Fish 10,000 Genomes Project
CNGB: China National Gene Bank
stLFR: single tube Long Fragment Read
NCBI: National Center for Biotechnology Information

## Data availability

The ten fish genome assemblies in the pilot have been deposited in the CNSA (https://db.cngb.org/cnsa/) of CNGBdb with accession codes CNP0000597 and CNP0000691.

## Author contributions

S-P. H., Q. Q., N-S. C., X.X., X. L., W. W. and G-Y. F. conceived and designed the study. L-D. Y., M-Q. Z., S.-S. L., and D.S., C. performed sample collection and sequencing. Y. S., S.-Y. Z., and X.-Y. H. performed assembly and annotation. X. L., Y. S., and G. F. wrote the manuscript. Y.-X. L., N.-S. C., and all other authors revised and read the manuscript.

## Acknowledgements

Thanks Dr. Inge Seim help us to polish the English writing of this manuscript.

---

**Supplementary Table 1.**
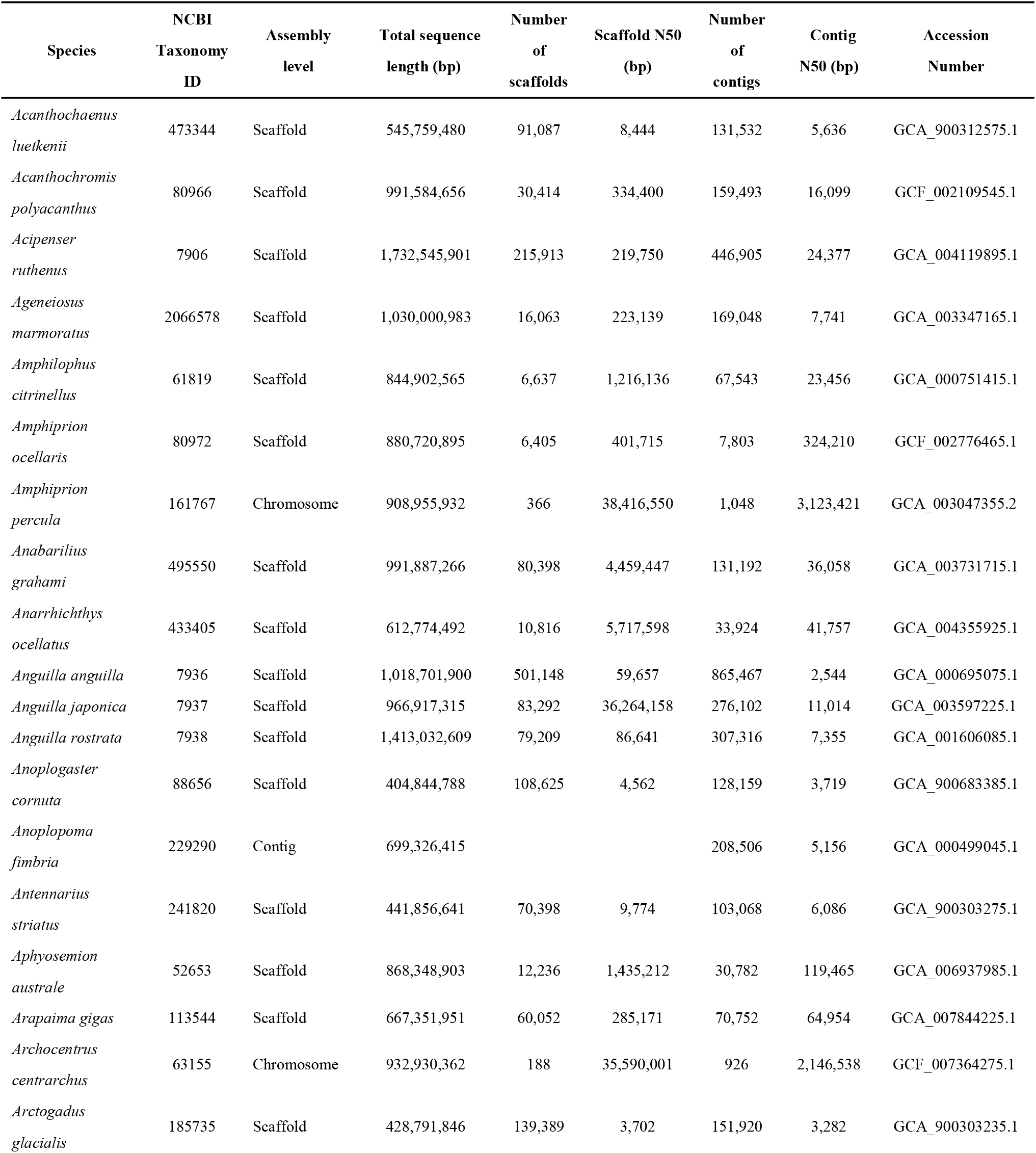
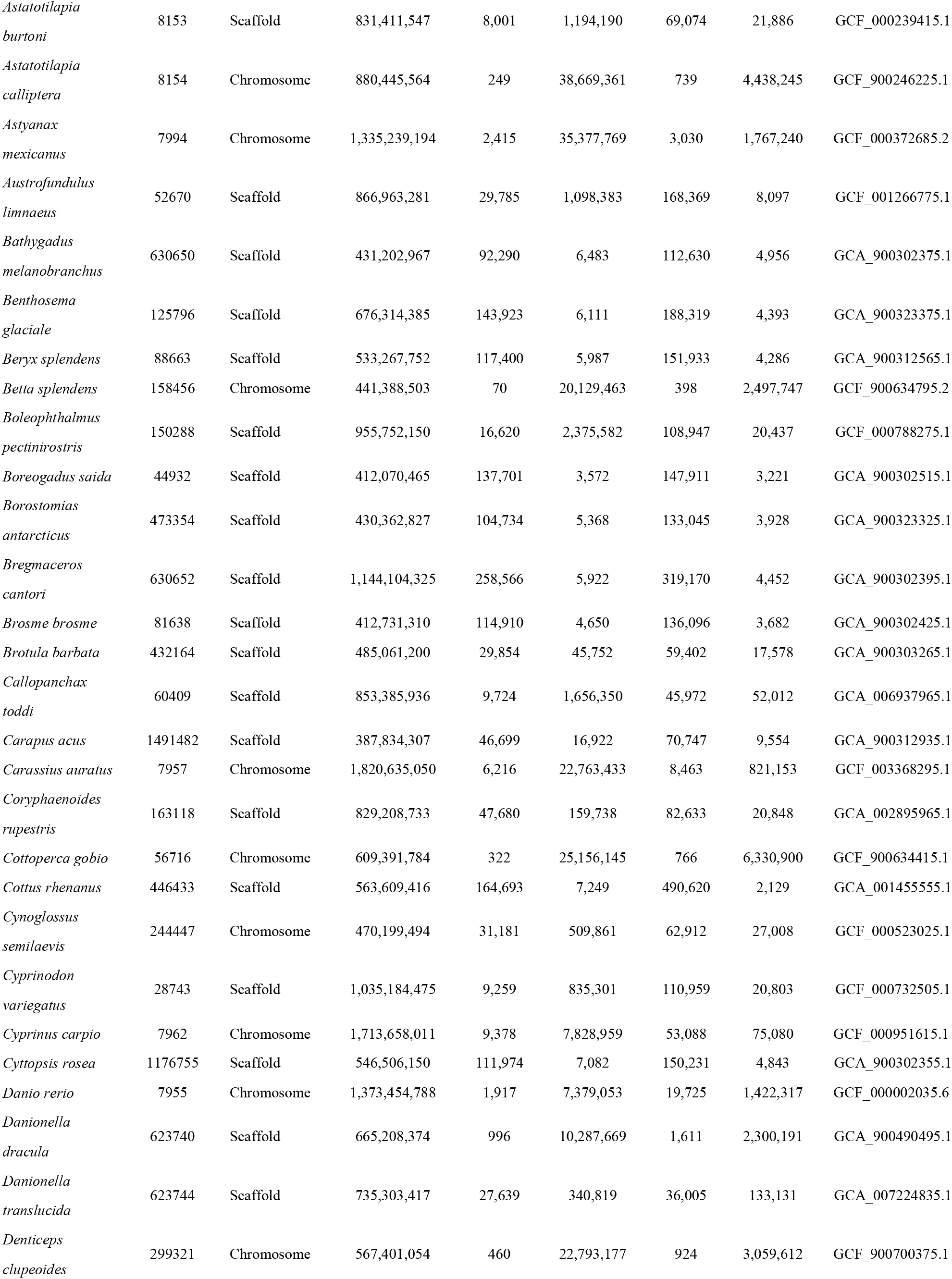
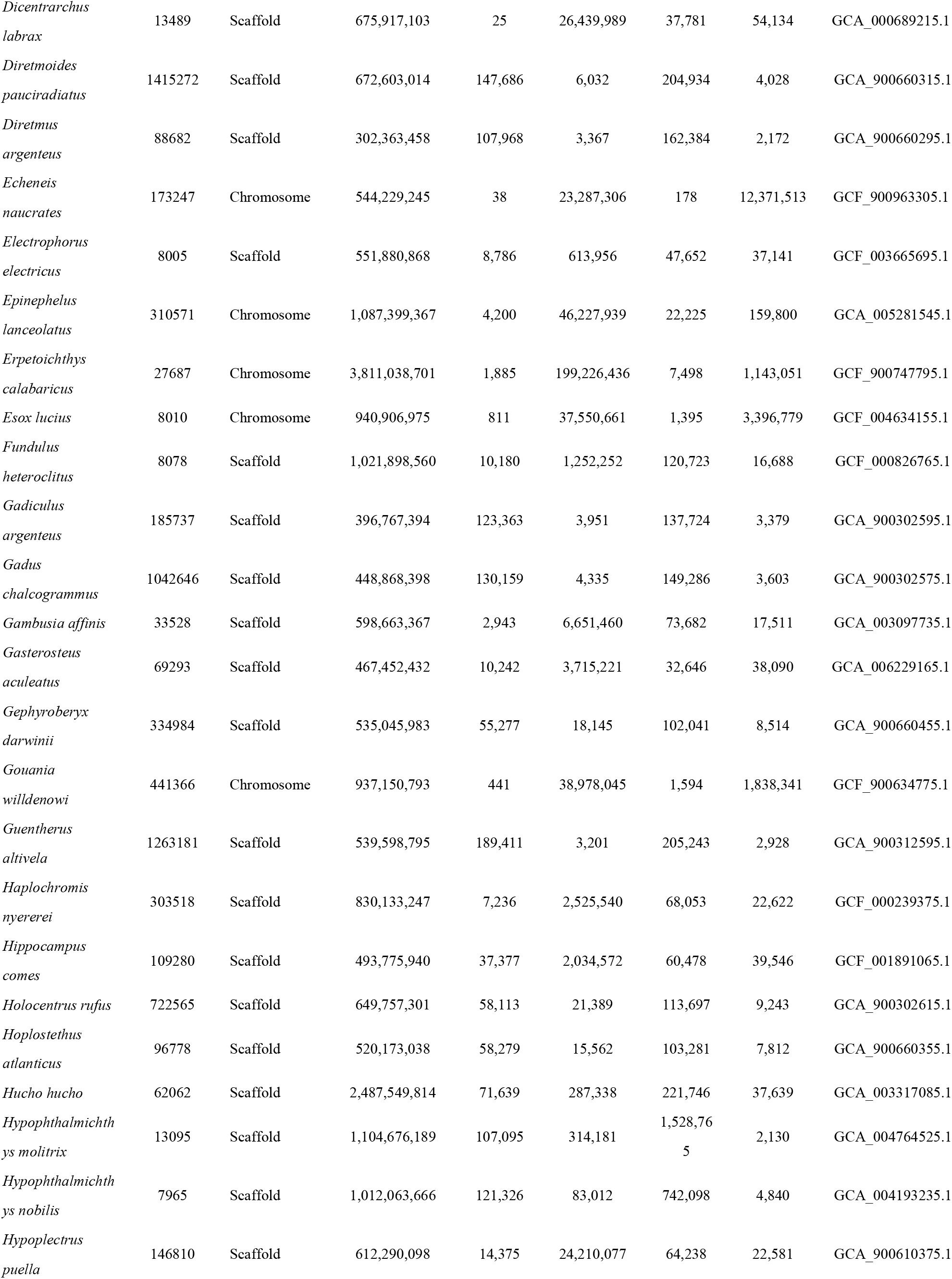
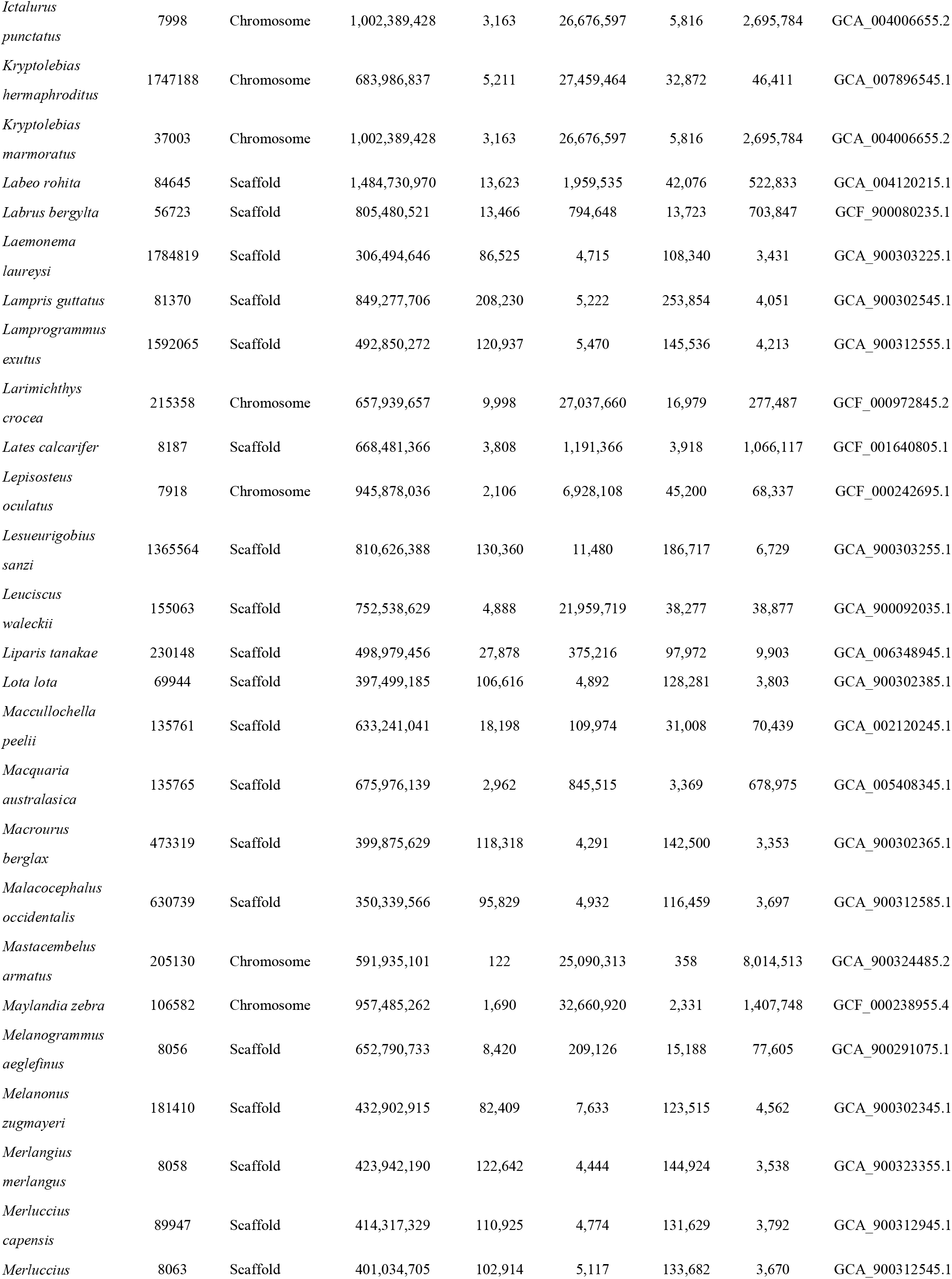
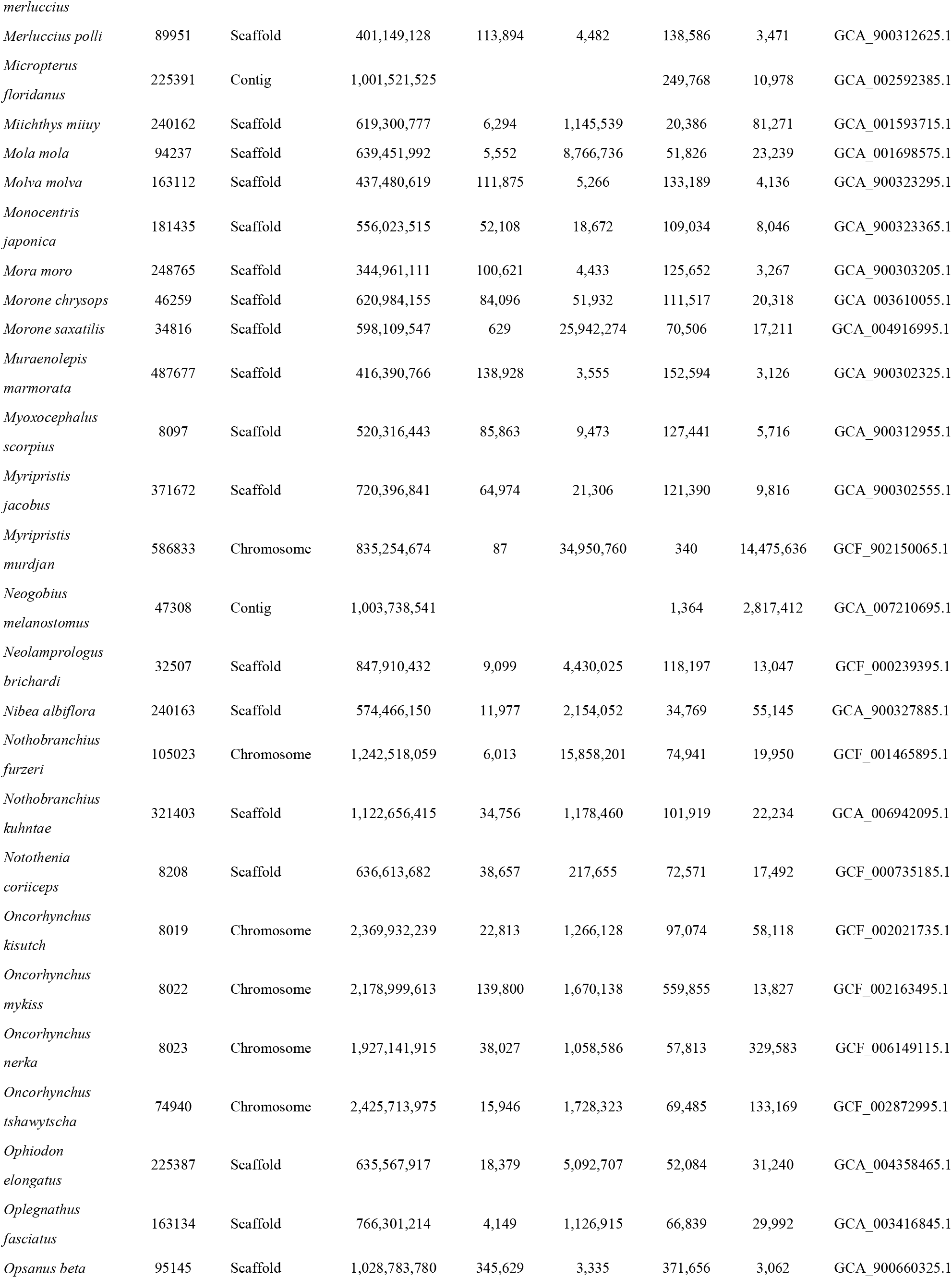
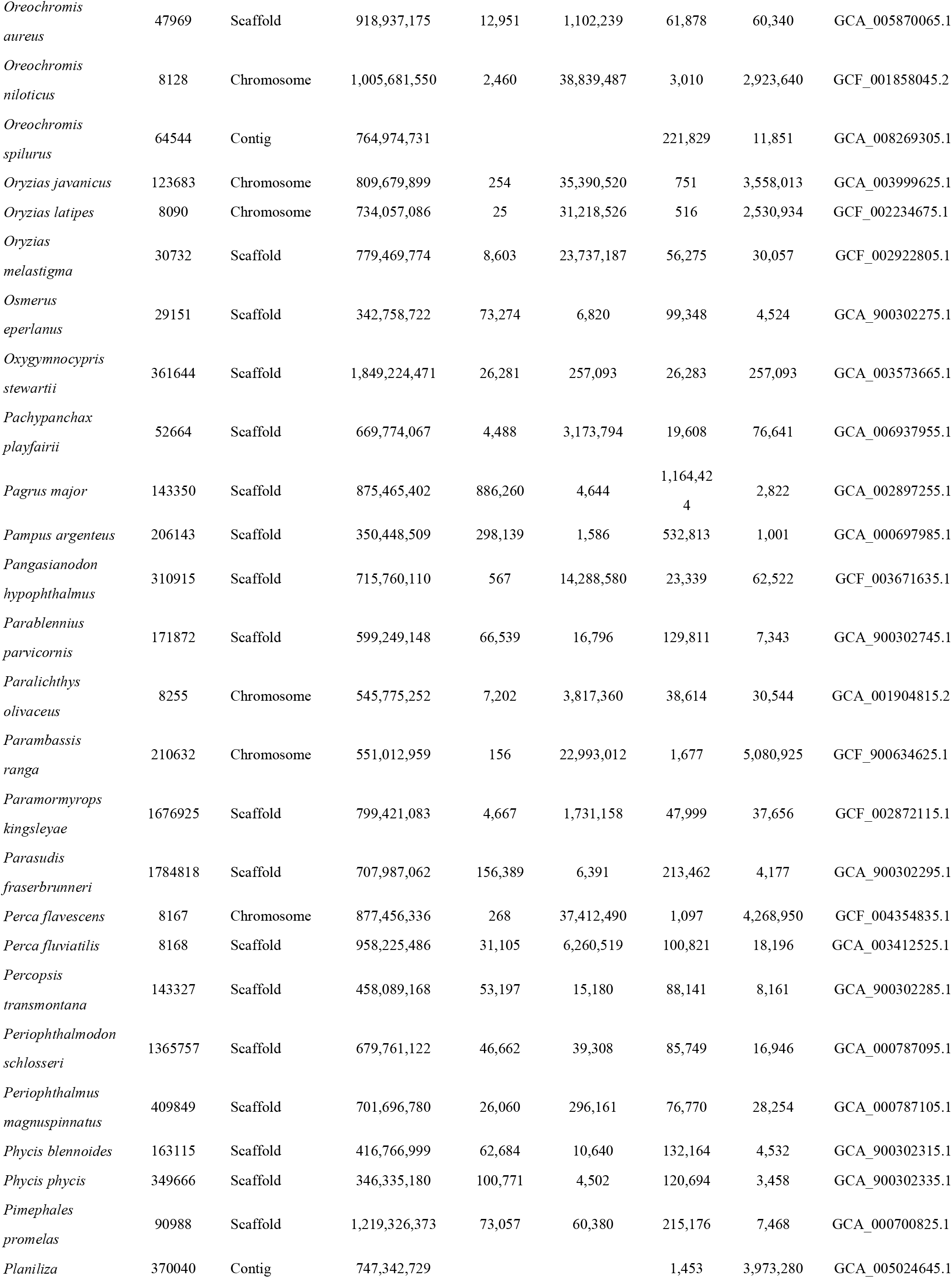
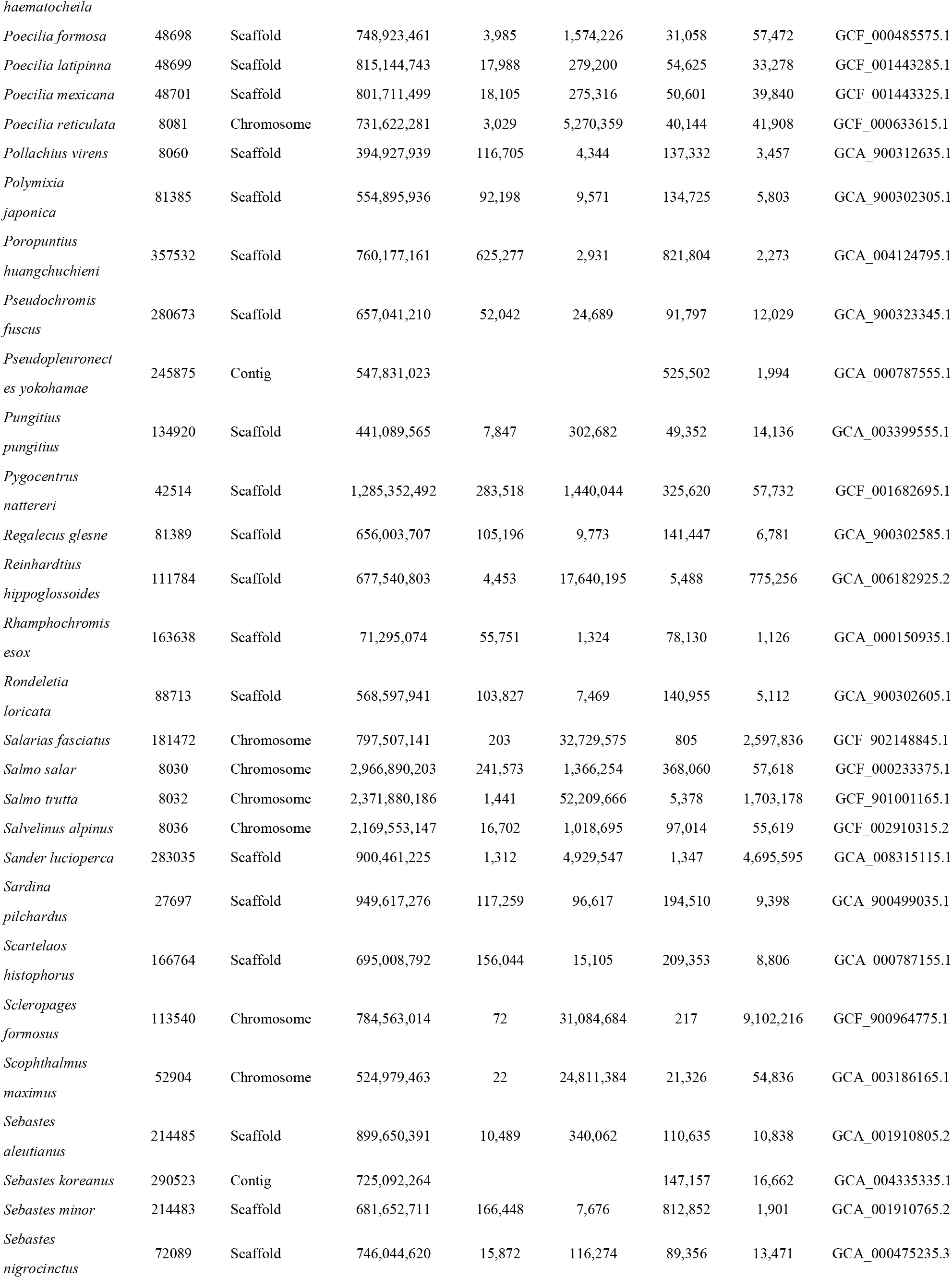
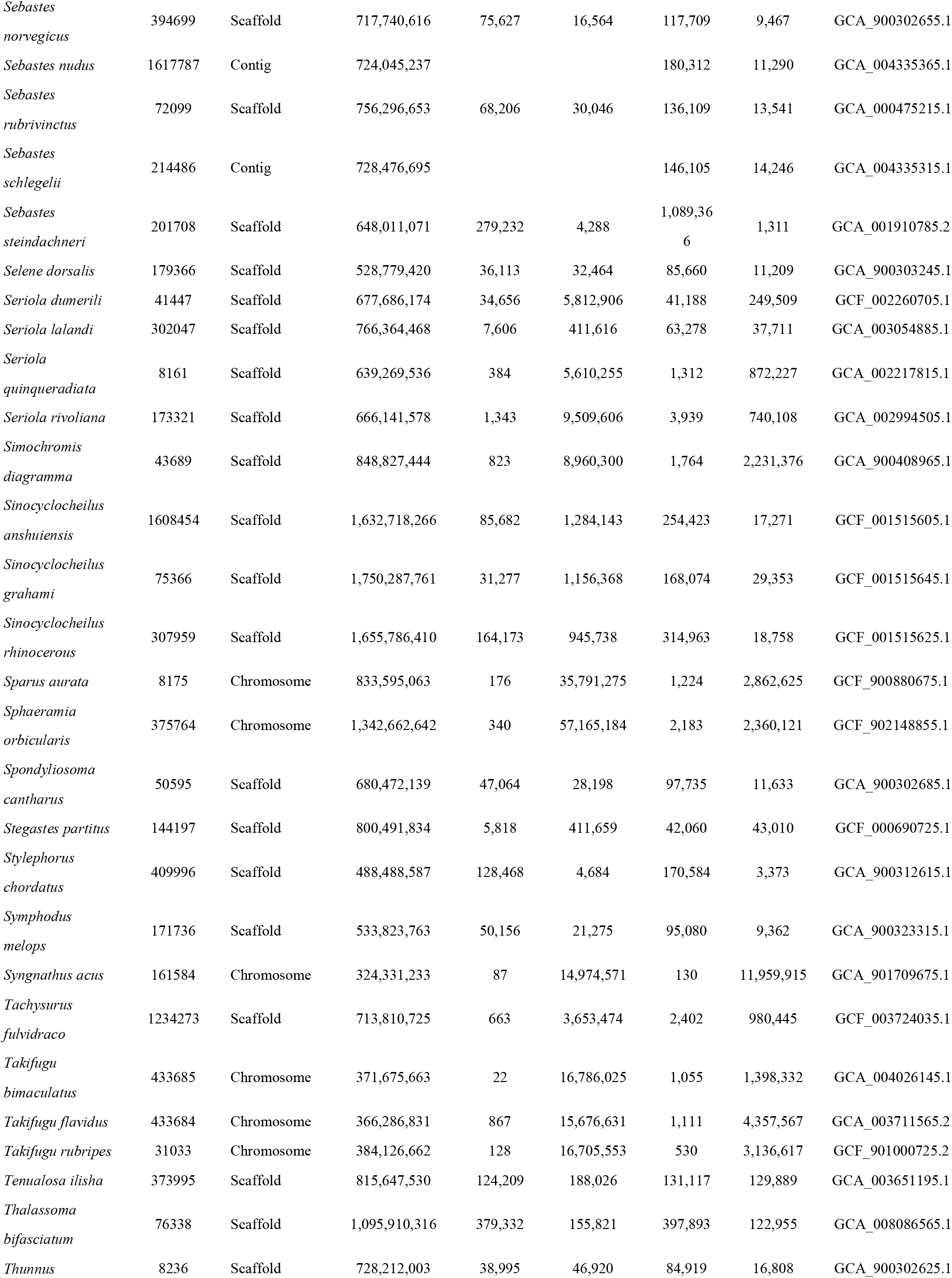
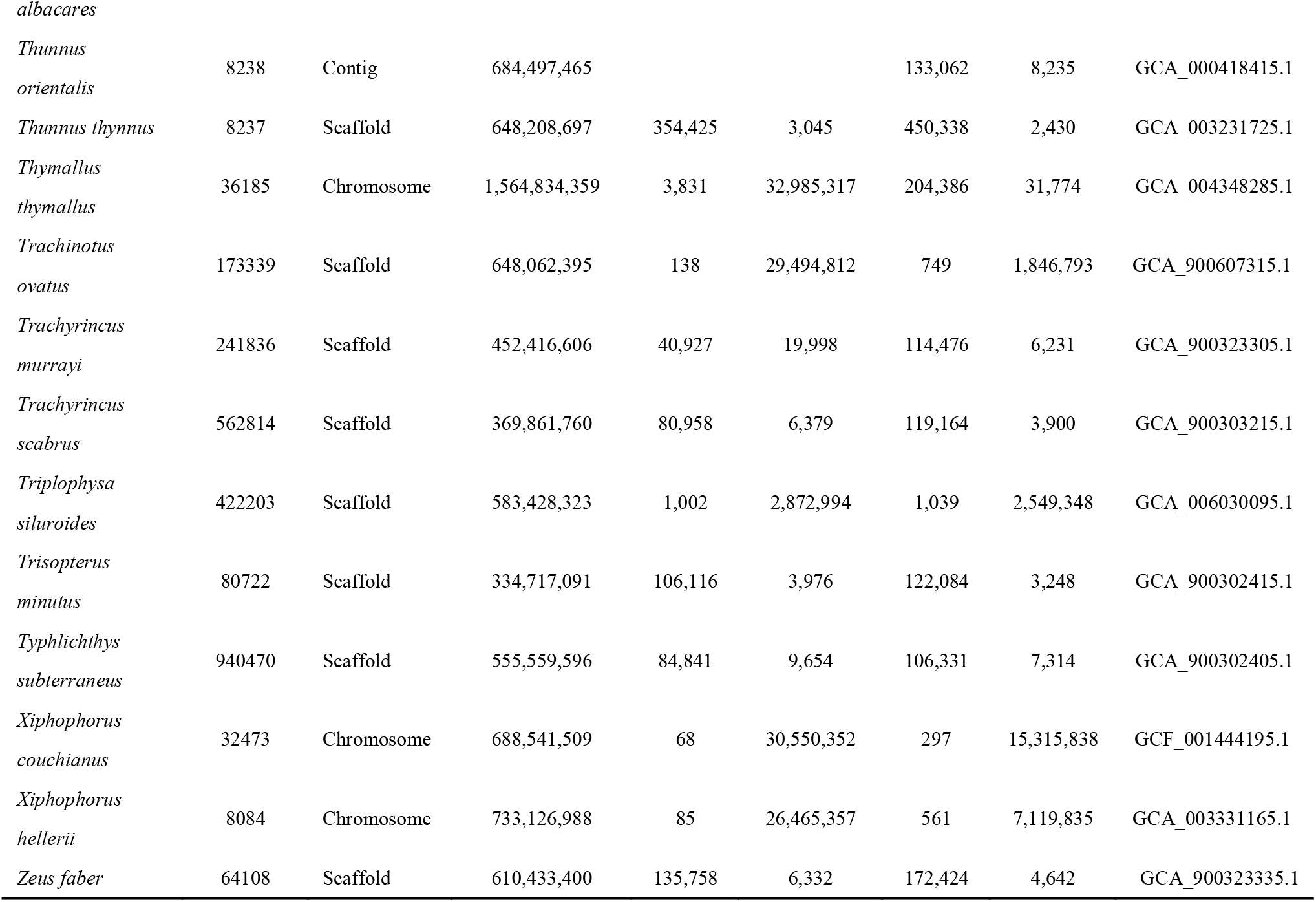
Assembly information of publicly available fish genomes. Extant fishes are divided into three major classes, among which Actinopterygii (ray-finned fishes) account for more than half.

**Supplementary Table 2.**
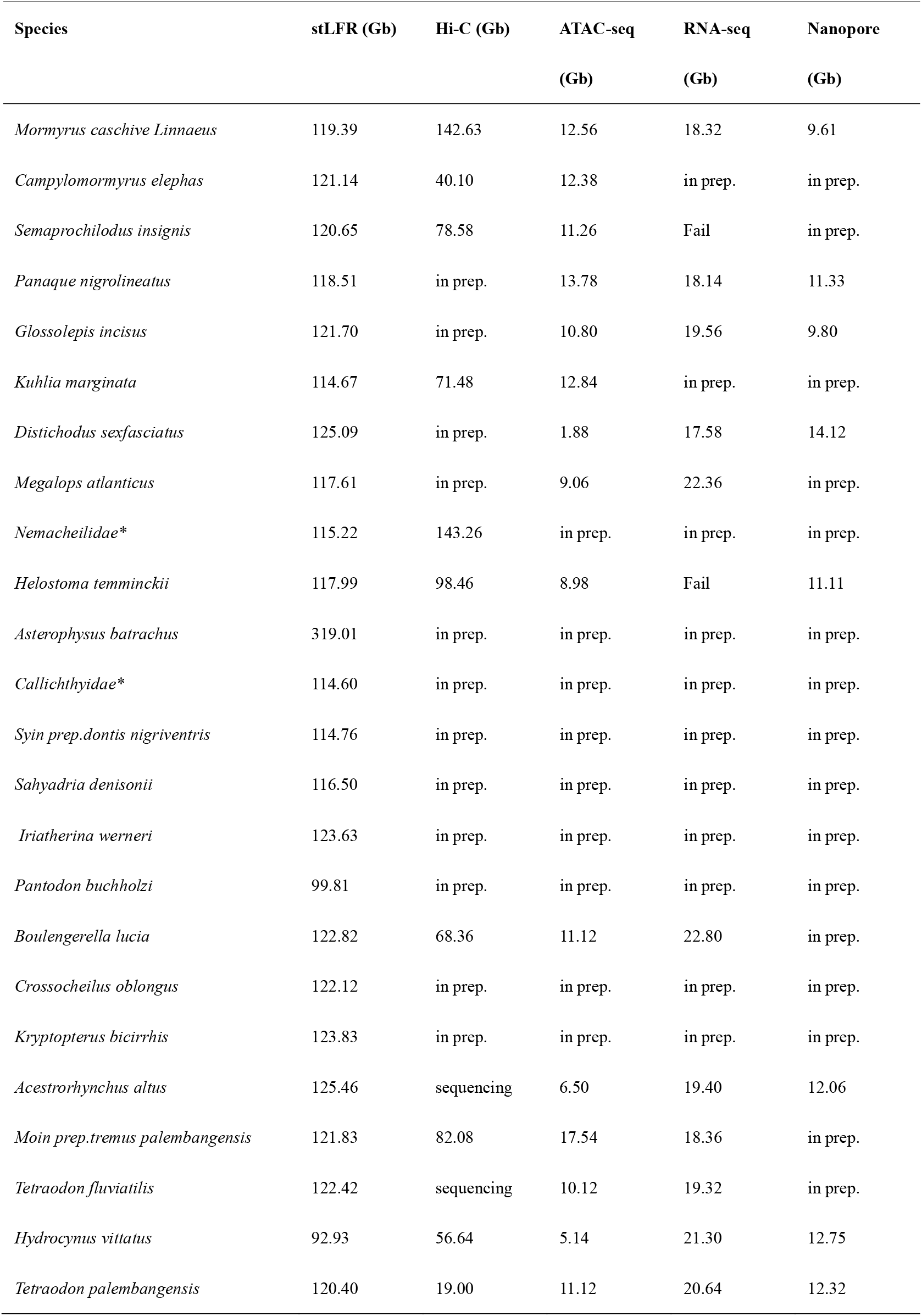
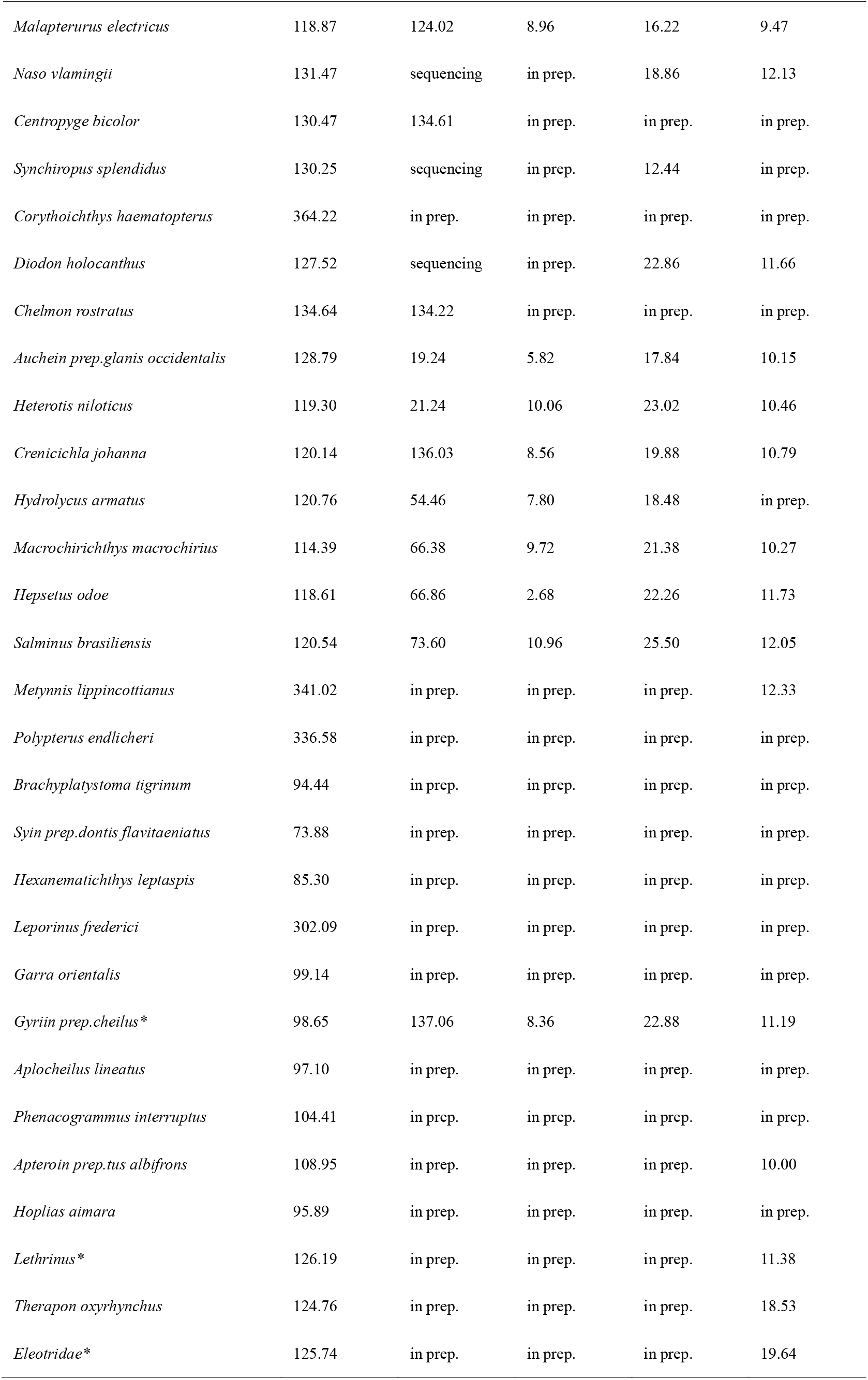
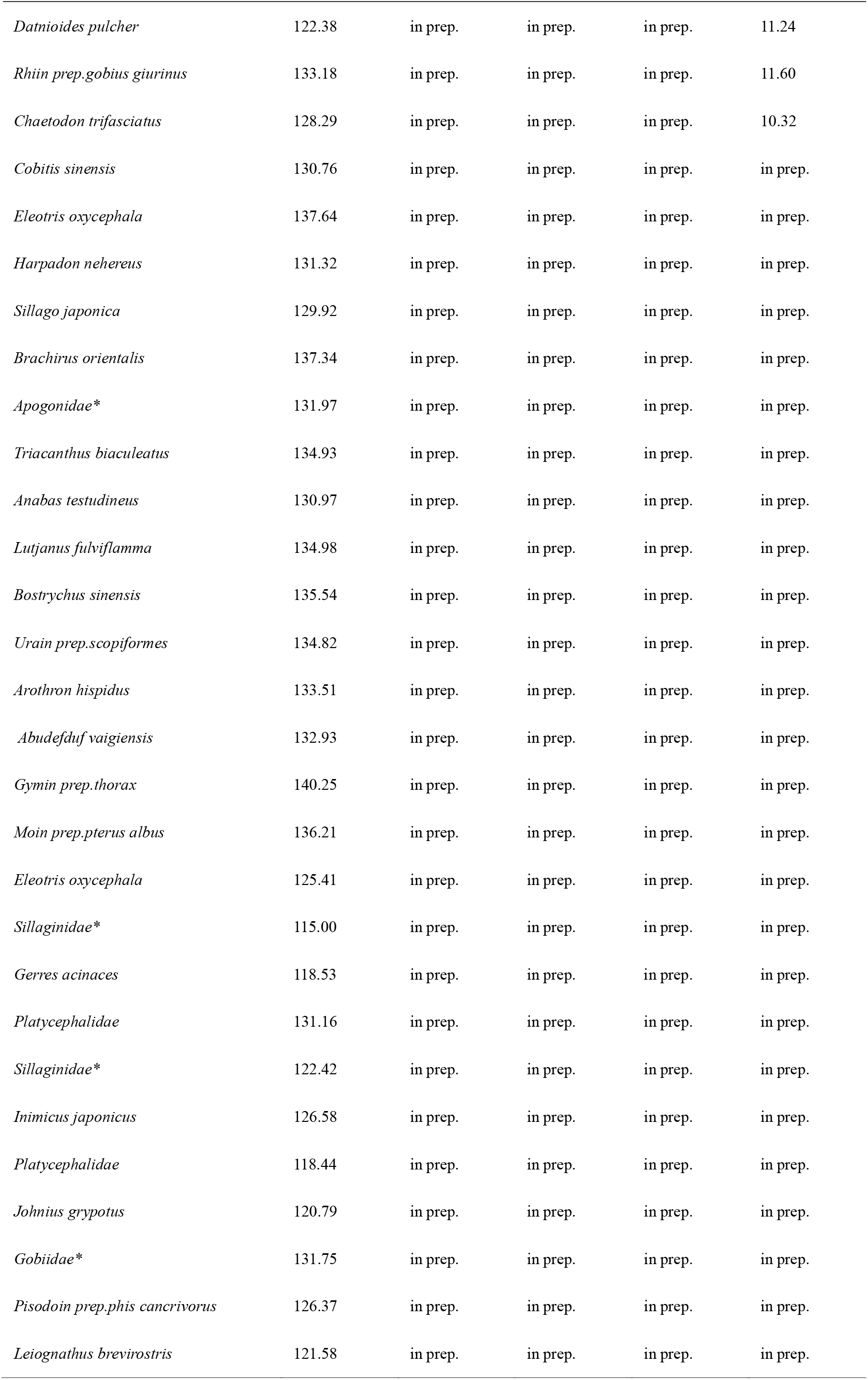
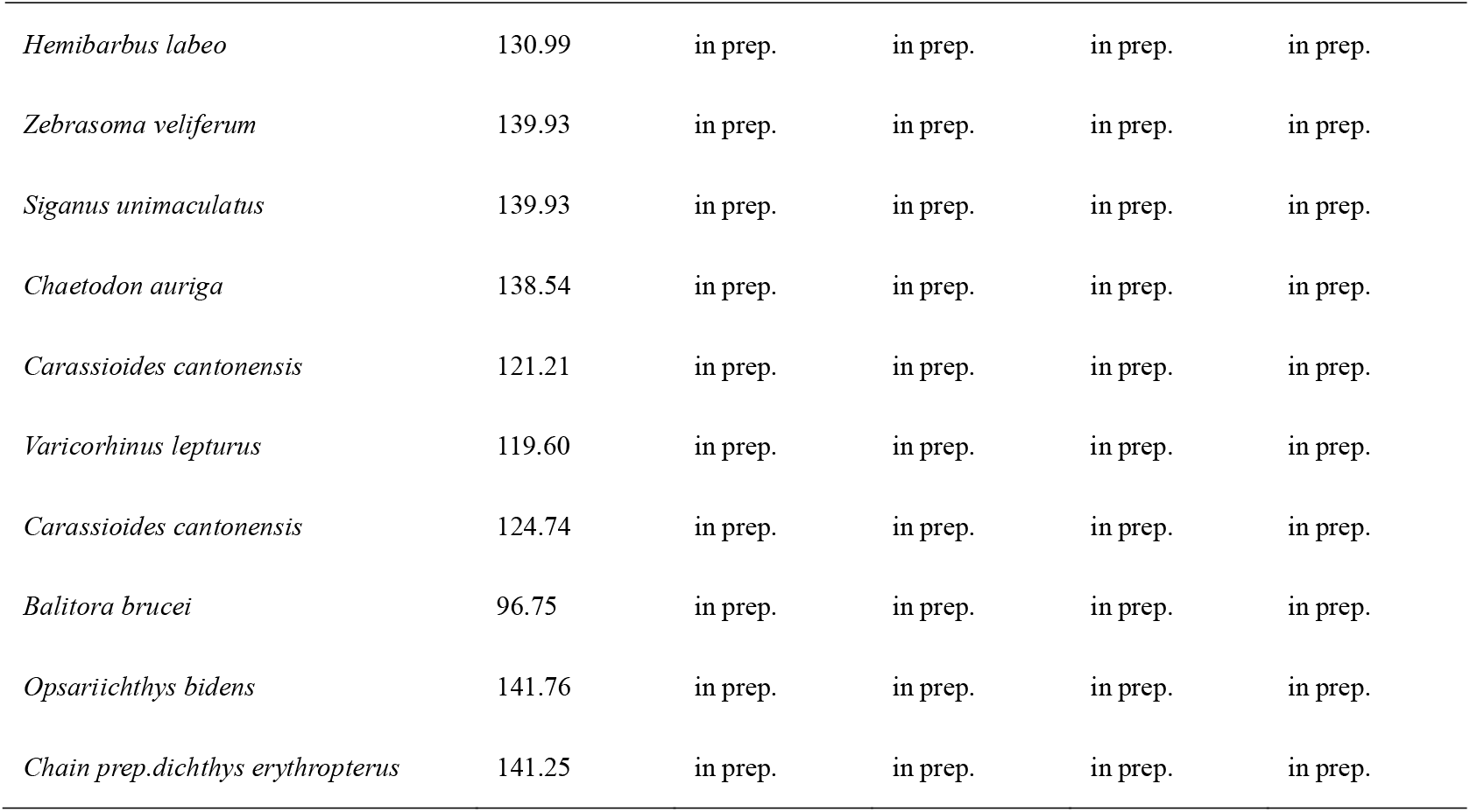
List of the 93 species sequenced in the Fish10K pilot study. An asterisk (*) denotes that a species could not be classified by morphological classification and need further evidence (i.e. genomics data) to be classified.

**Supplementary Table 3.** Assembly statistics of other 38 fish genome assemblies. An asterisk (*) denotes cartilaginous fish. A scaffold is a series of contigs that can be linked. N50 is the sequence length of the shortest contig (or contig) at 50% of the total genome length. BUSCO (Benchmarking Universal Single-Copy Orthologs) is a standard qualitative measure for assessing genome quality and completeness based on evolutionarily informed expectations of gene contents from near-universal single-copy orthologs. The highest BUSCO score is 100%. A higher BUSCO score indicates a better quality of the annotated genome.

| Strategy | Species | Assembly size<br>(bp) | Scaffold<br>N50 (bp) | Contig<br>N50 (bp) | BUSCO |
| --- | --- | --- | --- | --- | --- |
| I | *Raja porosa* * | 3,047,746,089 | 2,182,378 | 16,150 | --- |
|  | *Acestrorhynchus altus* | 955,799,133 | 9,632,287 | 48,769 | 94.0% |
|  | *Monotremus palembangensis* | 359,992,882 | 5,706,744 | 213,105 | 96.3% |
|  | *Tetraodon fluviatilis* | 326,757,569 | 7,010,509 | 47,398 | 96.0% |
|  | *Hydrocynus vittatus* | 1,007,380,147 | 3,263,334 | 25,092 | 88.7% |
|  | *Tetraodon palembangensis* | 359,651,202 | 7,038,803 | 218,260 | 96.2% |
|  | *Malapterurus electricus* | 798,763,704 | 3,588,439 | 48,402 | 90.3% |
|  | *Centropyge bicolor* | 681,516,296 | 4,420,410 | 115,524 | 96.2% |
|  | *Synchiropus splendidus* | 535,219,047 | 7,262,416 | 117,146 | 96.3% |
|  | *Auchenoglanis occidentalis* | 767,889,523 | 6,633,917 | 110,792 | 93.5% |
|  | *Crenicichla johanna* | 826,044,049 | 2,355,403 | 60,889 | 96.6% |
|  | *Hydrolycus armatus* | 806,097,948 | 3,692,170 | 20,392 | 86.0% |
|  | *Macrochirichthys*<br>*macrochirius* | 835,452,375 | 6,246,214 | 56,429 | 87.8% |
|  | *Hepsetus odoe* | 860,429,938 | 5,143,489 | 43,947 | 93.8% |
|  | *Salminus brasiliensis* | 942,321,778 | 2,364,359 | 39,978 | 89.3% |
|  | *Metynnis lippincottianus* | 1,347,978,869 | 1,154,248 | 21,525 | 89.1% |
|  | *Therapon oxyrhynchus* | 724,925,999 | 11,684,578 | 50,940 | 90.5% |
|  | *Rhinogobius giurinus* | 973,105,242 | 4,063,184 | 25,681 | 84.8% |
|  | *Mormyrus caschive* | 819,947,615 | 1,532,218 | 48,123 | 89.6% |
|  | *Linnaeus* |  |  |  |  |
|  | *Megalops atlanticus* | 1,033,524,629 | 7,259,301 | 65,702 | 95.4% |
|  | *Kuhlia marginata* | 674,579,433 | 4,757,558 | 40,209 | 92.8% |
|  | *Semaprochilodus insignis* | 1,178,289,156 | 1,780,909 | 29,765 | 80.9% |
|  | *Distichodus sexfasciatus* | 814,422,444 | 1,048,871 | 31,524 | 89.4% |
|  | *Glossolepis incisus* | 889,156,884 | 1,422,316 | 32,279 | 91.0% |
|  | *Campylomormyrus elephas* | 905,010,418 | 1,613,581 | 30,544 | 92.5% |
| II | *Parabramis pekinensis* | 984,227,144 | --- | 5,486,023 | 95.90 |
|  | *Pseudolaubuca sinensis* | 806,127,291 | --- | 11,561,046 | 96.00 |
|  | *Redfin culter* | 1,102,622,456 | --- | 2,126,716 | 94.60 |
|  | *Toxabramis swinhonis* | 1,038,263,990 | --- | 1,754,830 | 93.70 |
|  | *Squaliobarbus curriculus* | 1,066,071,259 | --- | 1,664,874 | 95.40 |
|  | *Ochetobius elongatus* | 929,258,157 | --- | 3,217,609 | 95.90 |
|  | *Opsariichthys bidens* | 841,061,991 | --- | 5,688,910 | 94.60 |
|  | *Nicholsicypris normalis* | 854,847,741 | --- | 2,288,299 | 95.50 |
|  | *Yaoshanicus arcus* | 835,190,039 | --- | 5,992,616 | 96.40 |
|  | *Xenocypris argentea* | 1,033,647,724 | --- | 5,841,742 | 96.30 |
|  | *Gymnocypris scolistomus* | 969,320,259 | --- | 1,907,468 | 92.00 |

**Supplementary Table 4.** Statistics of the existing fish species. Based on data from FishBase web site (accessed 10 September 2019) and “Fishes of the world” (5th ed.). Actinopterygii belong to Osteichthyes, whose number of species accounts for most all fishes.

| Type | Class | Order | Family | Species |
| --- | --- | --- | --- | --- |
| Chondrichthyes | Elasmobranchii | 12 | 51 | 1,190 |
|  | Holocephali | 1 | 3 | 56 |
| Osteichthyes | Actinopterygii | 67 | 475 | 32,869 |
| Total |  | 80 | 529 | 34,115 |

**Supplementary Table 5.** An Excel file with all the species list of Phase I.

## References

1. Venkatesh B, Lee AP, Ravi V, Maurya AK, Lian MM, Swann JB, et al. Elephant shark genome provides unique insights into gnathostome evolution. 2014;505 7482:174.

2. Wang K, Shen Y, Yang Y, Gan X, Liu G, Hu K, et al. Morphology and genome of a snailfish from the Mariana Trench provide insights into deep-sea adaptation. 2019;3 5:823.

3. Chen S, Zhang G, Shao C, Huang Q, Liu G, Zhang P, et al. Whole-genome sequence of a flatfish provides insights into ZW sex chromosome evolution and adaptation to a benthic lifestyle. Nature genetics. 2014;46 3:253.

4. Koepfli K-P, Paten B, Scientists GKCo and O’Brien SJJARAB. The Genome 10K Project: a way forward. 2015;3 1:57–111.

5. Vertebrate Genomes Project (VGP). https://genome10k.soe.ucsc.edu/vertebrate-genomes-project/. Accessed 10 September 2019.

6. Zhang GJN. Genomics: Bird sequencing project takes off. 2015;522 7554:34.

7. Zhang G, Li C, Li Q, Li B, Larkin DM, Lee C, et al. Comparative genomics reveals insights into avian genome evolution and adaptation. 2014;346 6215:1311–20.

8. BAT 1K. https://bat1k.ucd.ie/. Accessed 27 September 2019.

9. Cheng S, Melkonian M, Smith SA, Brockington S, Archibald JM, Delaux P-M, et al. 10KP: A phylodiverse genome sequencing plan. 2018;7 3:giy013.

10. DNA ZOO. https://www.dnazoo.org/. Accessed 27 September 2019.

11. Ostrander EA, Wang G-D, Larson G, vonHoldt BM, Davis BW, Jagannathan V, et al. Dog10K: An international sequencing effort to advance studies of canine domestication, phenotypes, and health. 2019.

12. Sun Y, Huang Y, Li X, Baldwin CC, Zhou Z, Yan Z, et al. Fish-T1K (Transcriptomes of 1,000 Fishes) Project: large-scale transcriptome data for fish evolution studies. 2016;5 1:18.

13. Wang O, Chin R, Cheng X, Wu MKY, Mao Q, Tang J, et al. Efficient and unique cobarcoding of second-generation sequencing reads from long DNA molecules enabling cost-effective and accurate sequencing, haplotyping, and de novo assembly. Genome research. 2019;29 5:798–808.

14. Waterhouse RM, Seppey M, Simao FA, Manni M, Ioannidis P, Klioutchnikov G, et al. BUSCO applications from quality assessments to gene prediction and phylogenomics. Mol Biol Evol. 2017;35 3 doi:10.1093/molbev/msx319.

15. FishBase. https://www.fishbase.se/search.php. Accessed 10 September 2019.

16. Nelson JS, Grande TC and Wilson MV. Fishes of the World. John Wiley & Sons; 2016.

17. Wenger AM, Peluso P, Rowell WJ, Chang P-C, Hall RJ, Concepcion GT, et al. Highly-accurate long-read sequencing improves variant detection and assembly of a human genome. 2019:519025.

18. The Fort Lauderdale Agreement. Reaffirmation and Extension of NHGRI Rapid Data Release Policies: Large-scale Sequencing and Other Community Resource Projects. Available at: https://www.genome.gov/10506537/reaffirmation-and-extension-of-nhgri-rapid-data-release-policies. Accessed 10 September 2019.

19. Birney E, Hudson TJ, Green ED, Gunter C, Eddy S, Rogers J, et al. Prepublication data sharing. 2009;461 7261:168–70.

